# Perinatal Oxycodone Exposure Causes Long Term Sex-Dependent Changes in Sensory and Reward Processing in Adult Mice

**DOI:** 10.1101/2022.02.15.480568

**Authors:** Elena Minakova, Marwa O. Mikati, Manish K. Madasu, Sineadh M. Conway, Justin W. Baldwin, Raylynn G. Swift, Katherine B. McCullough, Joseph D. Dougherty, Susan E. Maloney, Ream Al-Hasani

## Abstract

*In utero* opioid exposure is associated with lower weight and a Neonatal Opioid Withdrawal Syndrome (NOWS) at birth, along with longer-term adverse neurodevelopmental outcomes and mood disorders. While NOWS is sometimes treated with continued opioids, clinical studies have not addressed if long-term neurobehavioral outcomes are worsened with continued postnatal exposure to opioids. In addition, pre-clinical studies comparing *in utero* only opioid exposure to continued post-natal opioid administration for withdrawal mitigation are lacking. Therefore, we implemented a rodent perinatal opioid exposure model of Oxycodone (Oxy) exposure for comparison of long-term consequences of Oxy exposure until birth (Short Oxy) to the impact of continued postnatal opioid exposure (Long Oxy) spanning gestation through birth and lactation. Short Oxy exposure was associated with a sex-specific increase in weight gain trajectory in adult male mice. Long Oxy exposure caused an increased weight gain trajectory in adult males, sex-dependent changes in morphine conditioned place preference, and alterations in nociceptive processing in females. Importantly, there was no evidence of long-term social behavioral deficits, anxiety, hyperactivity, or memory deficits following Short or Long Oxy exposure. Our findings suggest that offspring with prolonged opioid exposure experienced some long-term sequelae compared to pups with opioid cessation at birth. These results highlight the potential long-term consequences of opioid administration as a mitigation strategy for clinical NOWS symptomology and suggest alternatives should be explored.

## INTRODUCTION

Opioid abuse in the United States has risen to epidemic proportions, with the US Department of Health declaring a public health emergency in 2017. Concurrently, the prevalence of opioid use disorder during pregnancy more than quadrupled, from 1.5 to 6.7 per 1,000 deliveries^1,2^. Consequently, there has been a significant increase in neonates hospitalized for Neonatal Opioid Withdrawal Syndrome (NOWS), a constellation of withdrawal symptoms following opioid exposure during pregnancy affecting the nervous, gastrointestinal, and respiratory systems^3,4^. Moderate-severe NOWS is managed in the neonatal intensive care unit by opioid replacement therapy to alleviate withdrawal^3^. To date, clinical studies have not addressed whether long-term neurobehavioral outcomes are worsened with continued postnatal exposure to opioids and adjunct agents used for withdrawal management^5^.

Epidemiological evidence suggests *in utero* opioid exposure is associated with lower birth weight and adverse neurodevelopmental outcomes in childhood, including cognitive deficits, impaired language development, attention deficit hyperactivity disorder, increased risk for autism spectrum disorder, decreased social maturity, and aggression^4,6–9^. However, large epidemiological studies evaluating longer term behavioral outcomes have been difficult due to confounding variables including: genetic factors, quality of caregiving, continued parental substance abuse, and socioeconomic variables which can significantly affect outcomes^10^. In addition, clinical studies comparing *in utero* opioid exposure to continued postnatal opioid administration for withdrawal mitigation have been lacking. Consequently, the development and evaluation of perinatal opioid exposure would enable exploration of conserved biological mechanisms mediating behavioral or sensory deficits.

Therefore, we designed a model in which opioid exposure spans preconception through early offspring development^4^, thus modeling gestational opioid exposure as well as postnatal opioid administration. This model also enables us to better understand the biological impact of opioid exposure on long-term development in the absence of confounding factors present in clinical observational studies. Specifically, we implemented subcutaneous, continuous infusion of Oxycodone (Oxy) at different durations enabling comparison of the longterm consequences of Oxy exposure until birth (Short Oxy) to the impact of continued postnatal opioid exposure (Long Oxy) spanning gestation through birth and lactation. We previously used this model to evaluate early outcomes (birth to one month of age) and showed Short Oxy exposure altered spectrotemporal features of affective vocalizations and caused sex-based differences in weight gain trajectories in offspring^4^. Long Oxy exposure further impacted weight, communicative behavior, and sensorimotor reflexes. These previous findings indicate that pups with Long exposure had worse overall outcomes, which suggests development of alternatives to opioid treatment for clinical management of NOWS symptomology^4^. Here we extended this analysis to explore later outcomes, assessing the effects of Oxy exposure on long-term social, anxiety-related avoidance, sensorimotor, cognitive, sensory and reward behaviors in juvenile and adult animals.

## MATERIALS and METHODS

### Animals

#### Animal Ethics, Selection and Welfare

All procedures using mice were approved by the Washington University Institutional Care and Use Committee and conducted in accordance with the approved Animal Studies Protocol. C57BL/6J mice (JAX, #000664) were housed in individually ventilated translucent cages (36.2×17.1×13cm, Allentown) or standard cages (28.5×17.5×12 cm) with corncob bedding and *ad libitum* access to standard lab diet and water. Animals were kept at 12/12 hour light/dark cycle with 20-22°C room temperature and 50% relative humidity.

A total of 48 dams were pair-housed and randomly selected to receive Oxy or Vehicle (Veh) infusion. All females were first bred to an age-matched male at postnatal day (P)60 to provide maternal experience and reduce this confound on pup survival of experimental litters. Following weaning of the first litter, treatment dams underwent surgical subcutaneous pump placement at P95 followed by a one-week recovery period. To evaluate the behavioral impact of early opioid cessation and abstinence in the developing offspring, half of the dams received an infusion of Oxy or Veh (Short Oxy and Short Veh) during pregnancy only followed by abstinence from the drug after delivery. The remaining dams continued to receive Oxy or Veh via the subcutaneous pumps through lactation (Long Oxy or Long Veh) to evaluate prolonged postnatal opioid exposure on developing offspring. To control for litter effects, each cohort was populated by multiple, independent litters. Litters were culled to 6 pups at P0-2 to remove effects of litter size. Two independent cohorts each comprising 40 Short Veh mice, 40 Short Oxy mice, 40 Long Veh mice and 40 Long Oxy mice were used. See **Table 1** for age at testing for each cohort. All mice were weaned at P21 and group-housed by sex and drug-duration assignment. Experimenters were female and blinded to group designations during testing. Preliminary work revealed significant dehydration and weight loss following weaning from the dam in Oxy-exposed offspring at P21. Therefore, to optimize survival, all mice were provided with Hydrogel (ClearH20; Highland Village, TX) and Bio-Serv Nutro-Gel Diet Purifield Formula (Hanover Park, IL) from P21-P30.

**Table 1.**
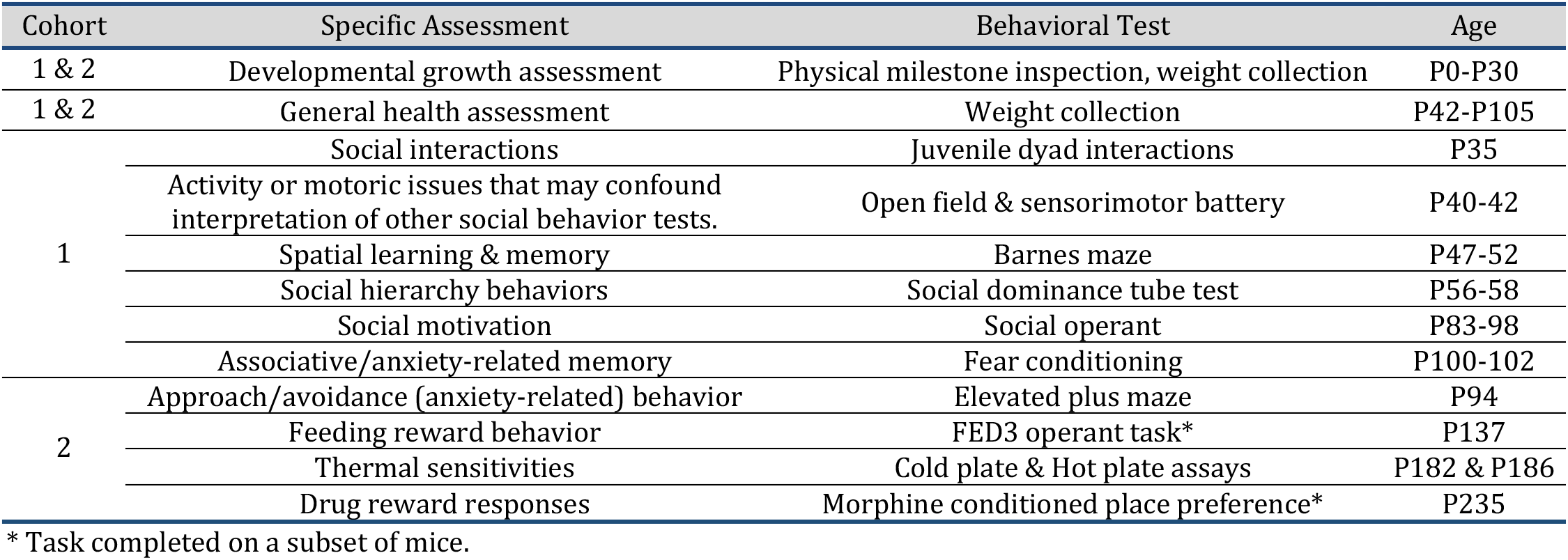
Order of and age at behavioral testing per cohort.

#### Drug Dosage, delivery system and surgery

Oxy (Sigma-Aldrich, Saint Louis, MO; Lot#: SLBX4974) dosage was as we previously reported^4^. Surgical procedures and drug delivery via Alzet pumps (Durect Corporation, Cupertino) were as previously detailed^4^. The Alzet pump was implanted and continuously infused either Oxy or sterile 0.9% NaCl (Veh) over a period of 4 weeks (0.23 μl/hr) or 6 weeks (0.15 μl/hr) for comparison of Short and Long Oxy and Veh, respectively.

### Behavioral Testing

Multiple behavioral assays (**Table 1**) across the same domain were employed to determine presence of behavioral, motor, or sensory disruptions. Cohorts were designed to ensure multiple stressful paradigms were not carried out on the same mice.

#### Developmental Growth Assessment

Mice were evaluated for achievement of physical and behavioral milestones from early development through adulthood. During the first two weeks postnatal, a visual inspection of physical milestone attainment was performed with evaluation for detached pinnae at P5 and eye opening at P14^4^. Weight, general health and appearance, were assessed at 14 time points: P0, P8, P21, P30, and weekly from 6-15 weeks.

#### Juvenile Dyadic Social Interaction

Juvenile social behaviors were assessed through juvenile dyadic interactions at P35 using a testing procedure performed as previously published with optimized lighting and software tracking method^11^. The test apparatus was as previously used with the addition of an LED light base (AGtEK Tracking Light Pad) underneath the surround to illuminate the mice from below (200 lux) and LED strip lighting (Commercial Electric) to illuminate the apparatus in low white light, allowing for optimal contrast and tracking accuracy. Tracking of both animals was completed in Ethovision XT v14 (Noldus) using the social interaction module. Time and frequency of head-to-head sniffs and anogenital sniffs exhibited by each the stimulus and test animal were analyzed. In addition, a social interaction outcome was calculated for the test animal by summing time spent engaged in head, body and anogenital area sniffing and body contact with the stimulus mouse.

#### Open Field Test

The open field test was used to assess the general activity and approach/avoidance (anxiety-like) behaviors of mice at P40 over a 1-hour period, based on previously published methods^12,13^. The test was performed using a transparent, acrylic testing apparatus (50×50×50 cm) enclosed within a white sound-attenuating box with dim red lighting at 10 lux.

#### Sensorimotor Battery

Balance, strength, and coordination were evaluated at P40-P42 by a battery of sensorimotor measures using previously published methodology^11^. The battery included assessment of walk initiation, basic balance (ledge and platform test), fine motor coordination (pole test), and strength (inclined and inverted screen tests).

#### Barnes Maze

The Barnes maze is a hippocampal-dependent spatial learning and memory task where animals learn the relationship between distal cues in the surrounding environment and a fixed escape location^14^. This task was conducted as previously described^12^ at P47-P52. Ethovision v14 (Noldus) was used to track the movement of the mouse within the apparatus and quantify distance traveled, frequency, latency, and duration of visits to the escape box or goal zone surrounding the escape hole, as well as total visits to other holes (errors). All males were tested first, followed by the females.

#### Tube Test of Social Dominance

Laboratory mice acquire social hierarchical rank behaviors between six-eight weeks of age which can be leveraged to examine social dominance behavior. This was performed at P56-P58 as previously described^11^.

#### Social Operant

Social motivation, including social reward seeking and social orienting was evaluated from P83-P98 using a social operant task. Procedures used here were as described previously^12^ (**Fig. S2C**) with minor alterations. Ethovision XT v14 (Noldus) was to track the movements of the test and stimulus mice during testing in their respective chambers. In addition, a white acrylic sheet was placed on the floor of the test chamber to optimize contrast and tracking accuracy. Custom Java tools and SPSS syntax were used to align the Ethovision tracking data with the timing of rewards in the Med Associates text data to extract presence or absence of each animal within the interaction zones during each second of every reward. For analyses, total hole pokes, correct and incorrect hole pokes, and rewards achieved were analyzed to assess learning, social reward seeking, and preference for active hole during fixed ratio 1 (FR1) training and fixed ratio 3 (FR3) testing. In addition, day to reach and number of mice that reached learning criteria to move on to FR3 were assessed. To examine motivation to elicit a social interaction, the breakpoint during progressive ratio (PR) 3 testing was analyzed.

#### Conditioned Fear Test

Fear conditioning was evaluated in our mice at P100-P102, following our previously described procedure^15^ (**Fig. 3D**). Shock sensitivity was evaluated following testing as previously described^15^.

#### Elevated Plus Maze

The elevated plus maze (EPM) test was used to assess negative affect and anxiety-like behavior of the mice at P94 based on previous methods^16,17^. Mice were habituated in the testing room, with dim lighting conditions at ~10 lux, for 2 hours prior to the experiment. Mice were then placed in an EPM. The maze is composed of two flat surfaces, equal in length and width of 75 cm, that form the shape of a plus sign. One arm has a vertical wall lining (closed arm), the other perpendicular arm does not have a wall lining (open arm). Each trial was 11 minutes with the exclusion of the first minute to account for any initial differences upon the placement of the mouse on the platform. The total time spent in either open arms, closed arms, or the center was measured and analyzed with Noldus EthoVision software. Additionally, distance traveled was calculated for each of the mice during the trials. The apparatus was cleaned out with 70% ethanol and water between each trial.

#### Feeding reward experimental devices (FED3) Operant Conditioning

To test for the acquisition of a feeding reward operant task, we trained the animals on the FED3 devices as previously described^18^. The animals began training at P137. Each animal was placed in a clean cage with isopad (BrainTree Scientific Inc.) bedding. The cage contained a FED3 device, and each mouse was trained on a FR1 task daily for 1-hour sessions. Following a nosepoke in the active port, a chocolate pellet reward (TestDiet) is released. The mice were trained until they met criterion, obtaining 80% correct nosepokes on three out of five training days. Mice were allowed 14 days to reach criterion (**Fig. S1A**). Following FR1, mice were trained on a FR3 schedule for three days where 3 correct nosepokes were rewarded with one chocolate pellet reward. After the 3-day FR3 schedule, the mice were put through an escalating PR task that started with FR1 then FR2 and increased by a factor of 2 after each correct round. The PR task was used to determine any differences in effort breakpoints between the groups of mice tested. The program was set to return to FR1 after 30 minutes of inactivity. During FR1 training, FR3, and the PR task, the number of active nosepokes, inactive nosepokes, and pellets earned were recorded for each mouse. Additionally, if the mice left any uneaten pellets, the number was also recorded to determine the number of pellets consumed.

#### Morphine Conditioned Place Preference (CPP)

The CPP test was used to assess changes in opioid-induced reward between mice. Mice at P235 were trained in a three-chamber, unbiased, and balanced conditioning apparatus as previously described^19^ (**Fig. 3A**). The apparatus had a horizontal context side, a vertical side, and a center without context. On the pre-test day, mice were allowed to access all three chambers of the apparatus for 30 minutes. Bias to a specific compartment was assessed based on the criteria of spending more than 67% of the time in one compartment during the pre-test as described^20^. Conditioning occurred over three consecutive days where each mouse received a saline injection in the morning (10 mL/kg, s.c.) and morphine (12.5 mg/kg s.c.) in the afternoon 4 hours after the morning session. Immediately after saline or morphine injection, the mice were placed in their assigned conditioning chambers for 30 minutes. To test for morphine place preference, the mice were allowed to explore all three chambers on the post-test day for 30-minute trial each. The preference scores were calculated as the difference in time spent in the drug associated compartment on the post-test day minus the pre-test day.

#### Hot/Cold Plate Assays

The hot/cold plate assay was used to determined changes in thermal pain thresholds between the mice as previously described^21^. Briefly, mice were habituated in the testing room for 30 minutes prior to the hot/cold plate assays. During the hot plate assay, individual mice were placed on a temperature controlled aluminum plate with the temperature set to 54.5°C for a maximum of 30 seconds to prevent tissue damage. A camera recorded behavior for post-experimental analysis. The latency to nocifensive behaviors: paw flicking, paw removal, paw licking and jumping was recorded. During the cold plate assay, the temperature was set to 4°C and the mice were placed on the apparatus for 5 minutes. Each trial was recorded and the frequency of jumping behavior was recorded for each mouse. The temperature of the plate was confirmed using temperature strips (Telatemp) and a thermometer prior to each trial. The plate was cleaned with 70% ethanol and water between each trial.

### Statistical Analyses

SPSS (IBM, v.25), Prism, and R were used for statistical analysis. Data was screened for missing values, influential outliers, fit between distributions and the assumptions of normality and homogeneity of variance. Variables that violated assumptions of normality were square root transformed as applicable (conditioned fear), assessed with, or confirmed by non-parametric tests. ANOVA, repeated measures (rmANOVA), Kruskal-Wallis, Mann-Whitney, linear modeling, and χ^2^ goodness of fit tests were used to analyze the behavioral data, with main factors of drug group and sex. The Huynh-Feldt or Greenhouse-Geisser adjustment was used to protect against violations of sphericity, where appropriate. Significant interactions between factors are reported. For significant main effects of sex, findings are segregated by sex. Multiple pairwise comparisons were Bonferroni corrected, when appropriate. Tukey’s post hoc test was used. Probability value was *p*<0.05. We analyzed the FED3 operant conditioning data using linear mixed effect models (with random effects on individuals) as mice were tested repeatedly. When analyzing summaries of mice that reached the training criterion, we used logistic regression. We analyzed the CPP data using linear models, as each mouse was tested once. For both linear and mixed models, we performed backwards model selection, starting with all appropriate three- and two-way interactions between predictors as well as additive effects. We assessed term significance via stepwise model comparisons and likelihood ratio tests (deviance tests for logistic regression, likelihood ratio tests for mixed models, and F-tests for linear models). Models were fitted in R version 4.1.2. (R Core Development Team 2021) using the package Imer Test^22^ and the code can be found on this website: https://github.com/MarwaMikati/Minakova_Mikati_2022. Test statistics and analysis details are provided in **Tables S1, S2, and S3**. The datasets generated for this study are available upon reasonable request.

## RESULTS

### Oxycodone exposure did not influence general functioning of offspring

To understand if perinatal opioid exposure influenced the general functioning in our mice, we examined activity levels, anxiety-related avoidance behavior, and sensorimotor abilities. As assessed in the open field test, Oxy exposure did not impact locomotor activity, with equal distances traveled by all groups (**Fig. 1A**). There was also no significant evidence for anxiety-like behavior as all groups spent comparable time in the center (**Fig. 1B**). To examine anxiety-related avoidance behaviors more directly, we conducted the EPM. Again, we saw no significant differences in percent time spent in the open arms between groups (**Fig. 1C**) nor in distance traveled (**Fig. 1D**), concurrent with the open field results. To confirm comparable sensorimotor capabilities between groups, a sensorimotor battery was performed to evaluate walk initiation, balance, motor coordination, and strength. No significant differences were seen between groups (**Fig. 1E-H**). Taken together, these data indicate Oxy exposure did not influence the animals’ abilities to move or increase spatial avoidance behaviors. This also eliminates concern regarding any movement confounds in the interpretation of all other testing. Therefore, we proceed to evaluate the effects of Long and Short Oxy exposure on the later functioning of behavioral circuits associated with opioids, i.e., nociception and drug reward.

**Figure 1:**
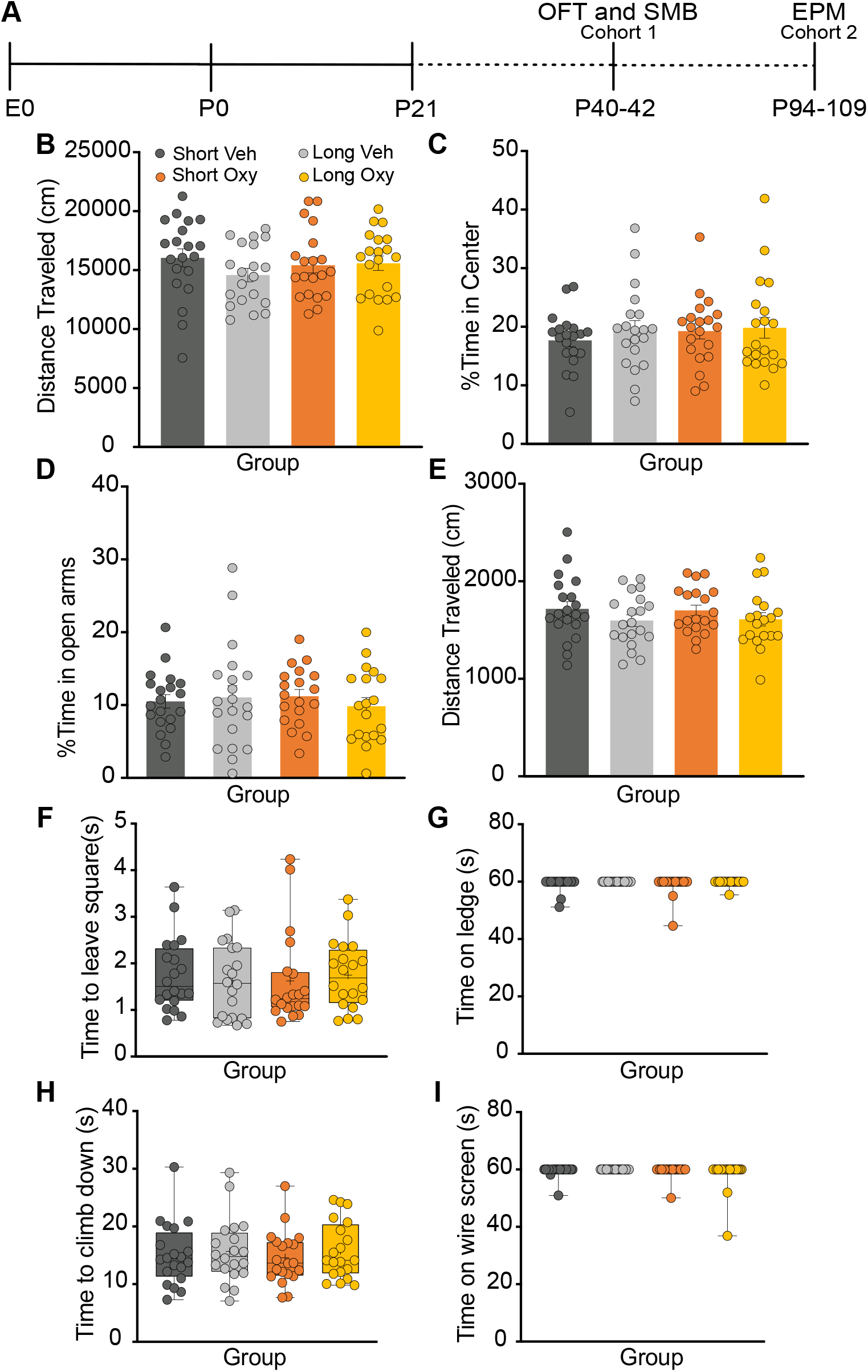
Oxycodone exposure does not impact anxiety-like behavior or sensorimotor development. **A.** Timeline of behavioral testing for the Open field test (OFT), Sensorimotor battery (SMB), and Elevated Plus Maze (EPM). **B.** The distance traveled in the open field test showed no significant differences in locomotion between groups. **C.** The percent of time spent in the center of the open field was comparable across groups **D-E.** In the EPM, percent of time spent in the open arms and total distance traveled were not different between groups. **F.** Motor initiation as measured by time to leave the center of a 20×20cm square surface was comparable between groups. **G.** Time able to balance on a ledge was comparable between groups. **H.** Latency to climb down a textured pole was not different between groups. **I.** Time to hang from a wire screen was not impacted by oxycodone exposure. Grouped data are presented as means ± SEM (n=80 for each assay) (**A-D**) and as boxplots with thick horizontal lines respective group medians, boxes 25th – 75th percentiles, and whiskers 1.5 x IQR (n=80 for each measurement) (**E-H**). Individual data points are filled circles.

### Oxycodone Exposure Increased Weight Gain Trajectory in Adult Male Mice

We evaluated the impact of Short and Long Oxy on litter size, appearance of physical milestones across the first two weeks postnatal and weight trajectory across the lifespan. There was no significant difference in litter size between Oxy or Veh exposed groups at birth (**Fig. 2A**). Weights were not significantly different between the Oxy groups and their respective Veh controls at birth (**Fig. 2B**), P8, P21, and P30 (**Fig. 2C**). Our previously published work revealed significant dehydration and weight loss following weaning from the dam in Oxy-exposed offspring from P21-P30^4^. Therefore, to optimize survival in our current study, all mice were provided with Hydrogel, Nutro-Gel Diet, and standard chow. This P21-P30 supplementation appeared to rescue the weight differences between Oxy and Veh groups, allowing us to study the long-term impact of exposure independent of any confounds due to a weight loss after weaning.

**Figure 2.**
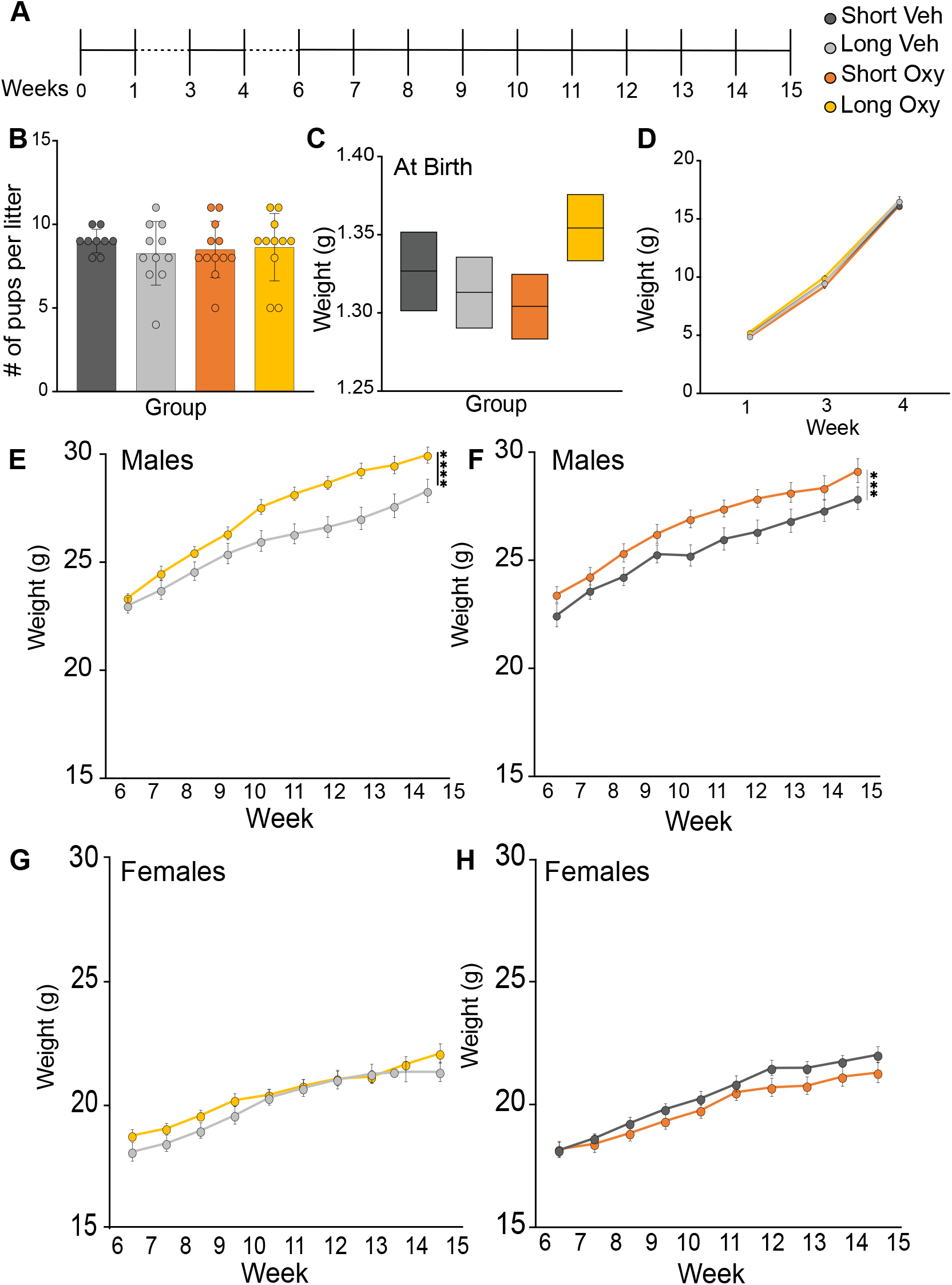
Oxycodone exposure resulted in increased weight gain in adulthood in males only. **A.** Timeline of weight measurements. **B.** Oxy-exposure in the dams did not impact litter size at birth. **C.** No differences in birth weight were observed for Oxy-exposed pups compared to Veh-exposed pups. **D.** Weight trajectories were similar for all groups from P8 – P30. **E.** Long Oxy exposure in males resulted in increased weights relative to Long Veh-exposed males in adulthood. **F.** To a lesser effect, Short Oxy exposure also increased weight gain in male mice compared to Short Veh-exposed male mice in adulthood. **G-H.** Neither Long (**G**) nor Short (**H**) Oxyexposure impacted female weight trajectories in adulthood. Grouped data are presented as means ± SEM with individual data points as filled circles, (n=160) for weight measurements. Statistical significance, ***p<.001, **p<.01, *p<.05.

We then evaluated the effects of Oxy exposure on long-term weight trajectory. Oxy exposure increased weight gain in male mice compared to Veh controls, particularly Long Oxy exposure, which was most pronounced in adulthood. Overall, male Long Oxy mice weighed more than males in either Veh group. This was particularly robust at weeks 10-15 (**Fig. 2D**). In addition, Short Oxy males weighed significantly more than Short Veh males at weeks 10-12 (**Fig. 2E**). Oxy exposure did not impact female weight trajectories (**Fig. 2F,G**). These data indicate that Oxy exposure resulted in a sex-specific increase in overall weight gain trajectory in adult males. Thus, exposure to Oxy during early development may potentially affect long-term weight gain and energy metabolism in males specifically.

Because opioids can regulate feeding behaviors through hedonic mechanisms, we investigated reward-based feeding behaviors using FED3 devices. Animals were trained on an operant conditioning FR1 task to receive a sugar pellet. Despite the difference in weight trajectory between the male oxy groups and their Vehicle controls, there was no difference in pellet number consumption trajectory by treatment group or sex, nor did the effect of treatment group vary by sex (**Fig. S1B; Table S2**). This suggests that there are no differences in consumptive behavior trajectories between these groups. Additionally, there were no significant differences between the groups in nosepoke accuracy over the FR1 training duration (**Fig. S1C; Table S2)**, and although the attainment of training criterion varied across sexes (80% accuracy for 3 out of 5 consecutive days; **Table S2**), Oxy exposure had no effect (**Fig. S1D; Table S2**). These findings suggest that Oxy exposure does not impact adult mice’s ability to learn the reward-based feeding task. Mice attaining criterion were subsequently tested on FR3 for three days followed by a PR test, neither of which showed differences between the groups (**Fig. S1E-F, Table S3**).

### Morphine conditioned place preference in adulthood shows sexual dimorphism

To investigate the effects of perinatal opioid exposure on drug reward behaviors in adulthood, we performed a morphine conditioned place preference test on a subset of the animals (**Fig. 3A**). We assessed the rewarding properties of morphine in both Short and Long Oxy groups and their respective controls. Interestingly, sex was a significant factor in the model, and the drugXduration interaction term had a *p*-value of 0.08 (**Table S1)**. This observation may be driven by the opposing trends in the Long Oxy male and female groups where the males show aversion to morphine at the 12.5mg/kg dose while females show a preference as compared to their respective Long Veh controls (**Fig. 3B,C**). This trend is not seen in the Short Oxy groups in males and females.

**Figure 3:**
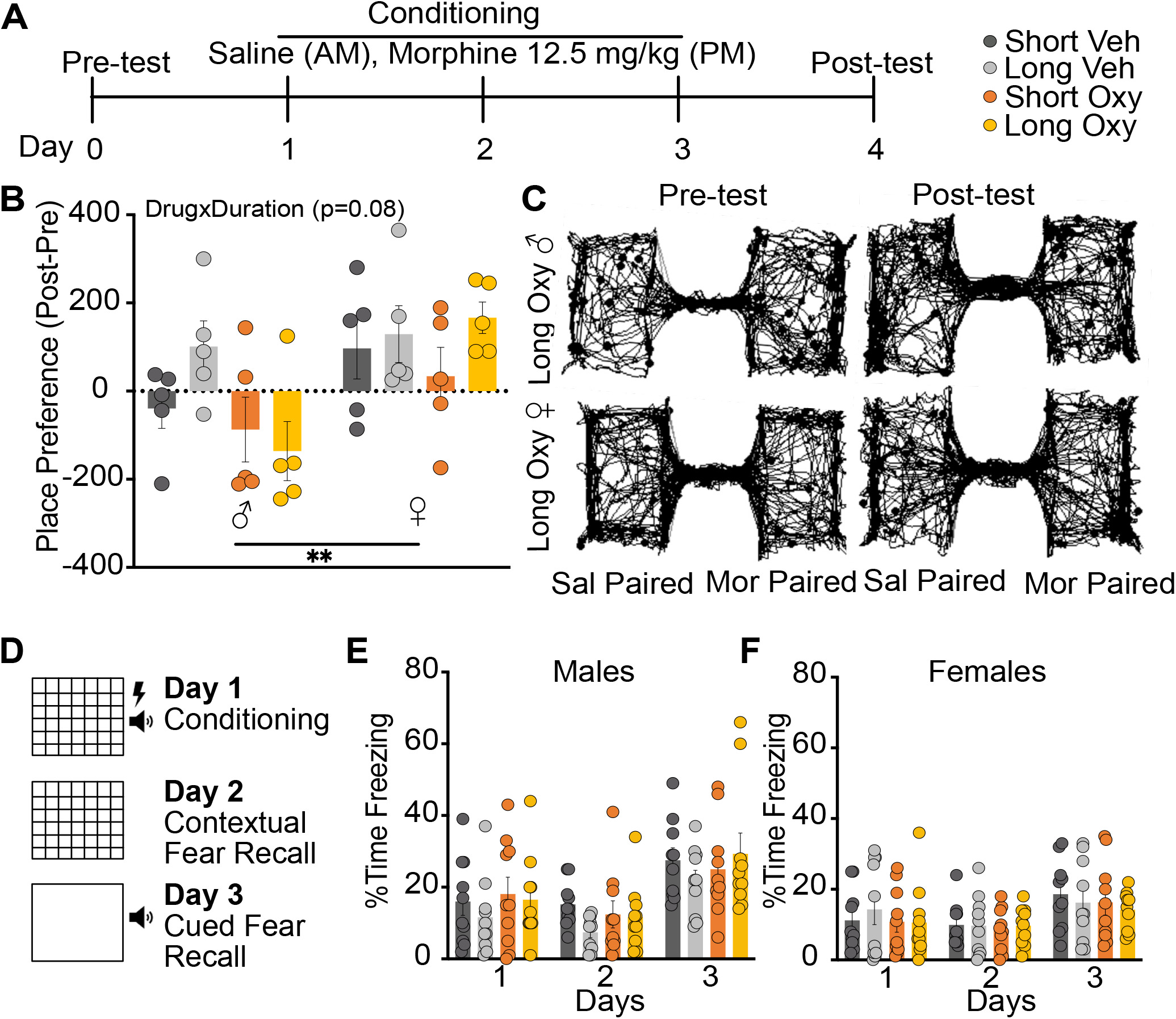
Oxy exposure leads to a sex-dependent trend in conditioning to opioid drug reward while fear conditioning remains the same. **A.** Morphine conditioning timeline and description. **B.** Morphine preference scores (n=40) reveal a sex difference and a trend in the DrugxDuration interaction term. **C.** Representative traces of a male mouse showing an aversion to the morphine paired chamber in the post test compared to the pretest while the female mouse shows a preference to the morphine paired chamber. **D.** Schematic describing the fear conditioning paradigm. **E.** No differences were found in the % of time spent freezing across the 3-day conditioning timeline in males **F.** or in females. Grouped data are presented as means ± SEM with individual data points as filled circles. Statistical significance, ***p<.001, **p<.01, *p<.05.

To understand if this sex-specific conditioning phenotype extended to aversive stimuli, we explored Oxy exposure on associative and anxiety-related memory in the fear conditioning paradigm (**Fig. 3D**). We did not observe differences between groups in the response to the pairing of tone/context with shock on day 1 (**Fig. 3E,F**), nor in sensitivity to shock current (**Table S4**). During conditioning test days 2 and 3, respectively, both Oxy-exposed groups displayed intact contextual and cued conditioning and no interaction with sex (**Fig. 3E,F**). These data suggest that the sex-specific responses induced by Oxy may be specific to drug reward.

The endogenous opioid system affects the motivational and rewarding aspects governing the development of juvenile social interactions and adult social behavior^23,24^. To evaluate the impact of Oxy on juvenile social behavior, we assessed the behaviors of our mice as juveniles in dyadic interactions. Oxy exposure, regardless of duration, did not alter juvenile social interactions. Specifically, frequency of or time spent sniffing (head or anogenital region) the social partner and body contact (**Table S5**) between the dyads were comparable between Oxy-exposed and control offspring. In addition, no differences were observed between groups for a composite score of total social exploration time (**Table S5**). To explore the impact of Oxy on mature social behaviors, we explored both hierarchical dominance and motivation to engage in social interaction. Mice display social hierarchies with dominant and submissive group members, which can be assessed using the tube test for social dominance^25^. The number of dominant and submissive displays were comparable between the Short Oxy and Short Veh groups, as well as the Long Oxy and Long Veh groups (**Fig. S2A,B**). In addition, no sex-specific effects on dominance behavior were observed amongst any treatment groups.

Since opioids can affect reward pathways influencing social interaction and behavior and to understand if Oxy may result in sex-specific responses to social reward, we assessed motivation for social interaction in our adult mice at P83 using our social operant task^12^ (**Fig. S2C**). The animals were trained in an operant conditioning task to receive a transient social interaction in response to a correct nose poke. No differences between groups were observed for nose poke accuracy (**Fig. S2D**) nor number of social rewards received (**Fig. S2E**) during FR1 training. Furthermore, Oxy exposure did not impact ability to reach learning criteria (**Fig. S2F,G**). We, again, did not observe differences among groups in rewards achieved when an increased effort was required during FR3 testing (**Fig. S2H**). Direct assessment of motivation to seek a transient social reward was performed through breakpoint testing using a PR3 reinforcement schedule. No group or group by sex differences were observed for breakpoint achieved (**Fig. S2I**). Together, our data suggest early Oxy exposure may sex-differentially affect later responses to opioids, yet leave social and aversive circuits intact.

Finally, to explore the impact of Oxy exposure on other learning and memory circuits, we examined spatial acquisition and reference memory performance in our mice using the Barnes maze. Neither Short or Long Oxy negatively impacted learning in this task as assessed by path lengths and latencies to the escape hole, or by total errors (non-target hole visits), across the five days of spatial acquisition (**Fig. S3A-C**). In addition, no group differences were observed for visits to or time spent in the goal zone surrounding the learned escape location during the probe trial (**Fig. S3D,E**), indicating intact spatial reference memory in the Oxy-exposed groups.

### Long Oxycodone exposure alters thermal sensation specifically in female mice

Since Oxy exhibits analgesic properties, we were interested in determining the effects of opioid exposure on thermal sensation (**Fig. 4A**). We observed that Long Oxy female mice were more sensitive to noxious cold, as shown by increased jumping, than Long Veh mice (**Fig. 4B**). Interestingly, we also saw a significant difference in the females between Long Oxy and Short Oxy groups. Similarly, we also investigated the effects of Oxy exposure on the hot plate test a noxious temperature of 54.5 °C. We found that in the females, the Long Oxy group showed an increase in latency to jump in comparison to the Long Veh group (**Fig. 4C**). This suggests that the Long Oxy females were less sensitive to the noxious hot temperature relative to the control group which may indicate an increased thermal pain tolerance. In conclusion, we show that Long Oxy females exhibited increased cold sensitivity and yet increased hot plate antinociception compared to the Long Veh controls. This contrasts with the male mice, which show no differences (**Fig. 4D, E**).

**Figure 4:**
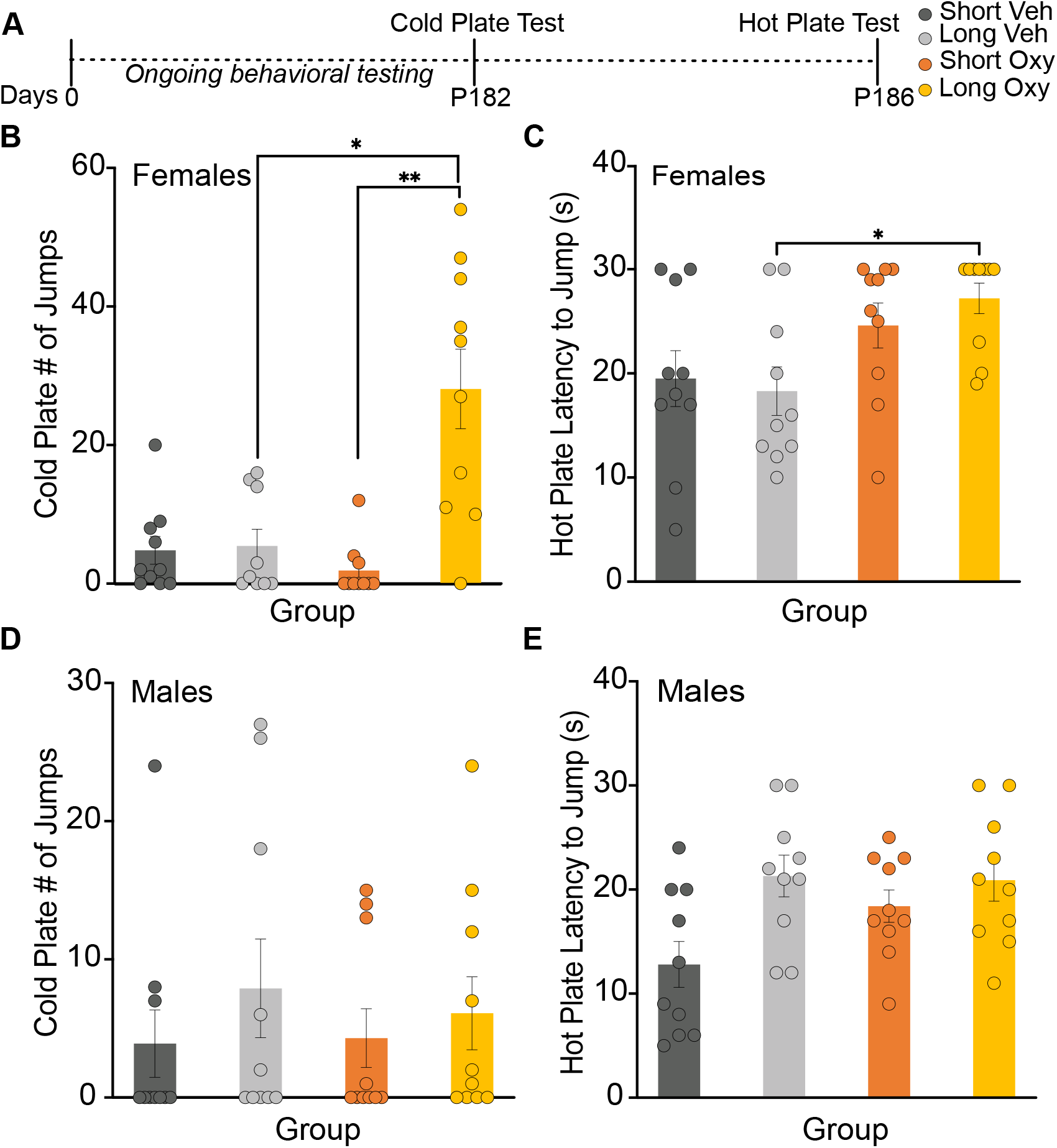
Long Oxy exposure alters thermal sensitivity specifically in female mice to both cold and hot temperatures. **A.** Timeline showing the ages at which the cold and hot plate assays were conducted. **B.** The number of jumps in the cold plate assay reveals that the female Long Oxy group is significantly higher than Short Oxy and Long Veh groups (n=40). **C.** The latency to jump in the hot plate assay shows that the female Long Oxy group has a significantly higher latency than the Long Veh group (n=40). **D-E** No differences were seen between the male groups in the cold plate number of jumps (n=40) (**D**) or the hot plate latency to jump (n=40) (**E**). Grouped data are presented as means ± SEM with individual data points as filled circles. Statistical significance, ***p<.001, **p<.01, *p<.05.

## DISCUSSION

Clinical studies have not addressed whether long-term neurobehavioral outcomes are worsened with continued postnatal exposure to opioids for withdrawal mitigation, or if pharmacotherapy should be limited^26^. Here, we present a murine model to investigate long-term neurobehavioral outcomes. We assess the impact of Short versus Long exposure to Oxy on long-term social, avoidance (anxiety-related) behavior, sensorimotor, cognitive, sensory and reward behaviors. Due to the role of opioids in the regulation of intrinsic natural reward behaviors and food intake^27^, we evaluated long-term weight trajectory and reward-mediated feeding behaviors in Oxy exposed mice. The literature on the effects of *in utero* opioid exposure on weight trajectory is limited and shows dependence on the specific opioid used^28,29^. Here, we demonstrated an overall increase in weight trajectory in both Short and Long Oxy male mice as compared to their Veh controls. However, we saw no differences in performance or pellet consumption between Oxy or Veh controls in our reward-mediated feeding task, suggesting the weight difference is not mediated by reward feeding. Yet, it is possible that we did not see differences in consumptive behavior due to the short one-hour task duration or the use of a sucrose-based reward. Future studies will be necessary to effectively evaluate comprehensive food consumption and indirect calorimetry as opioids have also been implicated in governing energy expenditure. For instance, a recent study has shown that Oxy exposed mice were less likely to engage in physical activity during the dark cycle using indirect calorimetry, and had higher body mass^30^.

In our investigation of changes in reward processing, we show that Long Oxy exposure differentially impacted CPP to morphine in adulthood between male and female mice. Long Oxy male mice developed an aversion to morphine while females show a similar preference to the Long Veh group. Based on prior literature of perinatal opioid exposure and CPP in adulthood^31,32^, we were not expecting to observe a sex difference. Therefore, we powered the study to test the drugXduration interaction while balancing for sex per NIH guidelines. Interestingly, our results correlate with a very recent study evaluating prenatal methadone exposure on drug-reward responses. The study demonstrated similar sex-dependent effects with males resistant to future alcohol reward while females were susceptible^33^. This aversion in adult male mice has also been observed in rodent studies of prenatal cocaine exposure^34^. Conversely, previous studies of prenatal morphine exposure have shown an enhancement in morphine CPP in adulthood^32,35^. The difference in drug-reward driven responses could be related to duration and onset of drug exposure during development^36^ as well as whether both males and females were tested in adulthood. Future studies will be necessary to further evaluate sex-specific behavioral effects of ontogenic Oxy exposure on morphine CPP and other non-opioid drugs of abuse and addiction.

There is a paucity of literature evaluating the long-term sensory effects of Oxy, a μ- and κ-agonist, following perinatal opioid exposure^37–40^. In our study, we demonstrate that Long Oxy females exhibit increased cold hypersensitivity and increased hot plate antinociception compared to the Long Veh controls. We see no differences in thermal sensitivity in the male mice tested. The altered sensory perception in the Long Oxy females is consistent with KOR activation, which has been implicated in enhancing cold sensitivity while also demonstrating antinociceptive properties toward noxious heat stimuli^21^. Our findings are the first to demonstrate sex-specific long-term sensory processing alterations following prolonged Oxy exposure in female offspring.

Here, perinatal opioid exposure does not appear to perturb circuitry governing social exploration, social reward behavior, anxiety-like behavior, or memory. Yet, our study suggests that offspring with prolonged opioid exposure experienced long-term changes affecting weight trajectory, sensory and reward processing, compared to pups with opioid cessation at birth. We included the Long Oxy exposure group to assess the consequences of continued opioid exposure after parturition, which is one of the mitigation strategies used in treating withdrawal in babies born to mothers with opioid use disorder. These results suggest judiciousness be applied to use of opioid administration as a mitigation strategy for clinical NOWS symptomology.

## ACKNOWLEDGEMENTS

We would like to thank Makenzie Norris, Justin Woods, and Victoria Collins for their assistance and support of this project.

## AUTHOR CONTRIBUTIONS

EM contributed to conceptualization, methodology, investigation, data curation, formal analysis writing (original draft preparation and editing), and funding acquisition for this project.

MOM contributed to methodology, investigation, validation, formal analysis, writing (original draft preparation and editing), and visualization for this project.

MKM contributed to methodology for this project.

SMC contributed to methodology for this project.

JWB contributed to the methodology and statistical analysis of this project.

RJS contributed to methodology for this project.

KBC contributed to methodology for this project.

JDD contributed to conceptualization, methodology, validation, resources, writing (draft editing), supervision, project administration, and funding acquisition for this project.

SEM contributed to conceptualization, methodology, data curation, resources, writing (draft editing), validation, supervision, project administration, and funding acquisition for this project.

RA contributed to conceptualization, methodology, validation, writing (draft editing), supervision, project administration, and funding acquisition for this project.

