## Supplementary Figures and Tables for "Perinatal Oxycodone Exposure Causes Long Term Sex-Dependent Changes in Sensory and Reward Processing in Adult Mice"

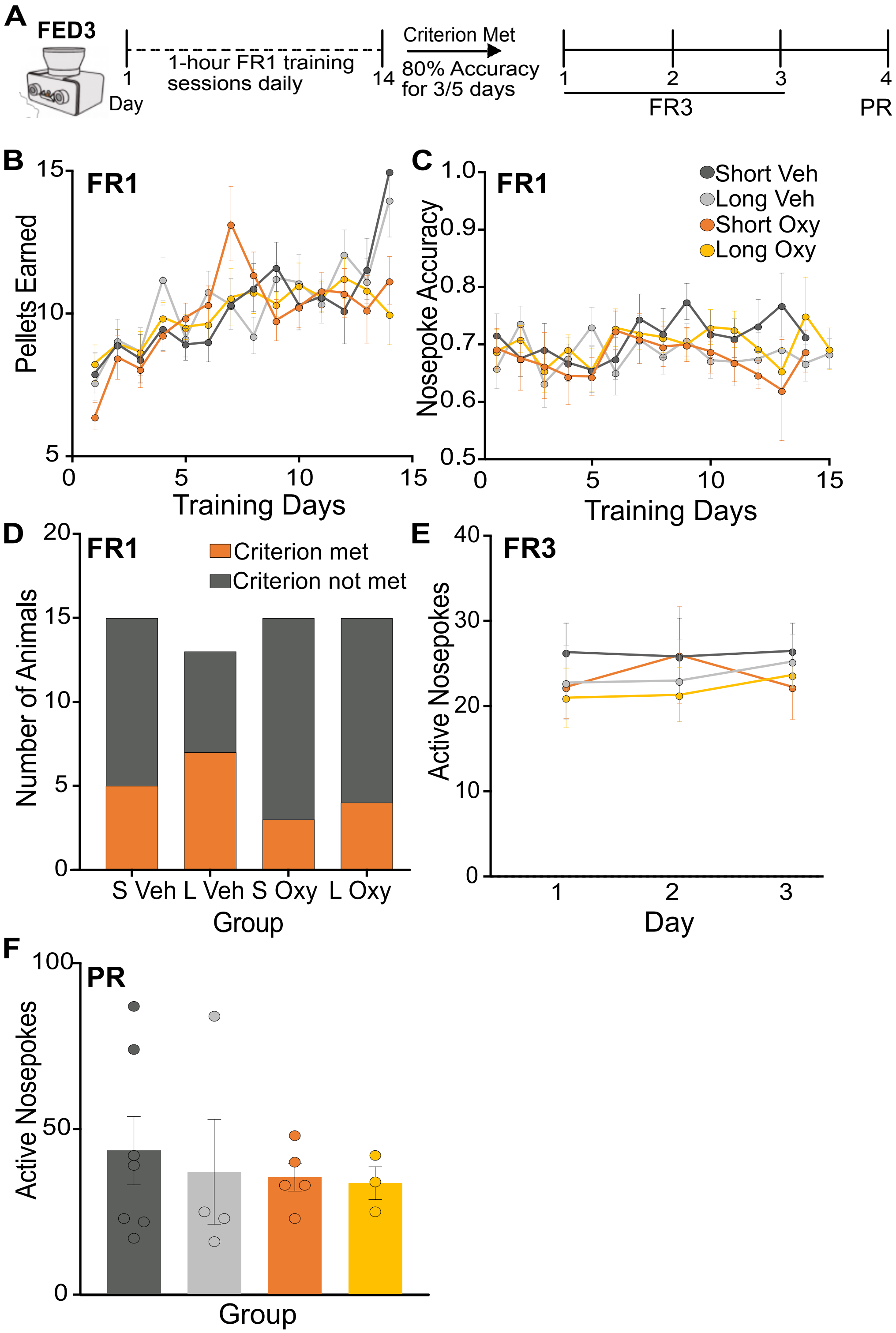


**Figure S1: Reward-based feeding behavior was not affected by Oxy-exposure. A.** Description of the FEDs operant task FR schedule and training criterion. **B**. The average number of pellets earned during daily FR1 training (n=58) **C.** Nosepoke accuracy as measured by correct/total nosepokes across FR1 training days were comparable across groups **D**. No differences were observed in the number of animals that attained the learning criterion across groups **E.** The number of nosepokes during the 3-day FR3 schedule for the animals that attained the training criterion showed no differences between groups. **F.** The number of active nosepokes during the progressive ratio test in the animals that met the training criterion reveal no differences between the groups.

Table S1. Statistical analysis results from experiments displayed in Figures 1-4.

| Figure | Variable | Statistical Test | Comparison | Output | *p* value |
|---|---|---|---|---|---|
| 1 | Open field distance traveled | ANOVA | Group | *F*(3,72)=.830 | .481 |
| | Open field time in center | ANOVA | Group | *F*(3,72)=.293 | .830 |
| | EPM % time in open arms | ANOVA | Group | *F*(3,70)=.292 | .831 |
| | EPM distance traveled | ANOVA | Group | *F*(3,70)=1.045 | .379 |
| | SMB Walk | Kruskal-Wallis | Group | *H*(3)=1.655 | .647 |
| | SMB Ledge | Kruskal-Wallis | Group | *H*(3)=2.402 | .493 |
| | SMB Pole | Kruskal-Wallis | Group | *H*(3)=.647 | .886 |
| | SMB wire hang | Kruskal-Wallis | Group | *H*(3)=2.275 | .517 |
| 2 | Litter size | ANOVA | Group | *F*(3,39)=.319 | .812 |
| | Pup weight, P0 | ANCOVA | Group | *F*(3,350)=1.824 | .142 |
| | Pup weight p8, P21, P30 | rmANOVA | Group | *F*(3,156)=1.939 | .126 |
| | | | Group*Age | *F*(3.6,188.4)=.310 | .853 |
| | Weight weeks 6-15 | rmANOVA | Group | *F*(3,151)=3.307 | .031 |
| | | | Group*Sex | *F*(3,151)=2.709 | .047 |
| | | simple main effect | Males, Group | *F*(3,151)=5.188 | .002 |
| | | | Group*Age | *F*(13.5,677.3)=1.071 | .381 |
| | | | Group*Age*Sex | *F*(13.5,677.3)=3.979 | .000002 |
| | | simple main effects | Males, week 8, Group | *F*(3,151)=3.098 | .029 |
| | | Kruskal-Wallis † | | *H*(3)=6.125 | .106 |
| | | simple main effects | Males, week 10, Group | *F*(3,151)=7.643 | .00009 |
| | | Kruskal-Wallis † | | *H*(3)=14.602 | .002 |
| | | simple main effects | Males, week 11, Group | *F*(3,151)=7.128 | .0002 |
| | | Kruskal-Wallis † | | *H*(3)=13.963 | .003 |
| | | simple main effects | Males, week 12, Group | *F*(3,151)=7.526 | .0001 |
| | | Kruskal-Wallis † | | *H*(3)=14.505 | .002 |
| | | simple main effects | Males, week 13, Group | *F*(3,151)=7.573 | .0001 |
| | | Kruskal-Wallis † | | *H*(3)=14.145 | .003 |
| | | simple main effects | Males, week 14, Group | *F*(3,151)=5.027 | .002 |
| | | Kruskal-Wallis † | | *H*(3)=10.484 | .015 |
| | | simple main effects | Males, week 15, Group | *F*(3,151)=4.328 | .006 |
| | | Kruskal-Wallis † | | *H*(3)=10.199 | .017 |
| 3 | Morphine conditioning preference | ANOVA | Drug | *F*(1,36)=0.236 | .630 |
| | | | Duration | *F*(1,36)=1.962 | .170 |
| | | | Sex | *F*(1,35)=10.894 | .002 |
| | | | Drug*Duration | *F*(1,33)=3.12 | .087 |
| | | | Drug*Sex | *F*(1,33)=2.719 | .109 |
| | | | Duration*Sex | *F*(1,33)=.173 | .680 |
| | | | Drug*Duration*Sex | *F*(1,32)=2.24 | .144 |
| | Cond Fear, Day 1 M3-5 | rmANOVA | Group | *F*(3,72)=.170 | .916 |
| | Cond Fear, Day 2 | rmANOVA | Group | *F*(3,72)=1.468 | .230 |
| | Cond Fear, Day 3 | rmANOVA | Group | *F*(3,72)=.654 | .641 |
| 4 | Cold plate jumps | Kruskal-Wallis | Group | *H*(3)=11.12 | .011‡ |
| | | Kruskal-Wallis | Male, Group | *H*(3)=1.619 | .655‡ |
| | | Kruskal-Wallis | Female, Group | *H*(3)=15.18 | .002‡ |
| | Hot plate paw withdrawal latency | Kruskal-Wallis | Group | *H*(3)=5.738 | .125‡ |
| | | Kruskal-Wallis | Male, Group | *H*(3)=10.58 | .014‡ |
| | | Kruskal-Wallis | Female, Group | *H*(3)=.039 | .998‡ |
| | Hot plate latency to jump | Kruskal-Wallis | Group | *H*(3)=10.61 | .014‡ |
| | | Kruskal-Wallis | Male, Group | *H*(3)=8.010 | .046‡ |
| | | Kruskal-Wallis | Female, Group | *H*(3)=9.527 | .023‡ |

‡Indicates Bonferroni corrected observed p value. †Indicates non-parametric confirmation of parametric results of non-normal data.

Table S2. Statistical details of the FED3 data analysis.

| Response | Model | Predictor | Chi squared | Df | P |
|---|---|---|---|---|---|
| Pellets earned | Linear mixed effects model | Group x Sex x day | 6.098 | 3 | 0.1069 |
| | | Group x Sex | 3.1284 | 3 | 0.3722 |
| | | Sex x day | 3.1648 | 1 | 0.07524 |
| | | Group x day | 2.1678 | 3 | 0.5383 |
| | | Sex | 10.669 | 1 | 0.001089 |
| | | Group | 0.3178 | 3 | 0.9567 |
| | | Day | 86.737 | 1 | <0.0001 |
| Nosepoke accuracy | Linear mixed effects model | Group x Sex x day | 2.9462 | 3 | 0.4 |
| | | Group x Sex | 7.5513 | 3 | 0.05625 |
| | | Sex x day | 0.4883 | 1 | 0.4847 |
| | | Group x day | 3.0691 | 3 | 0.3811 |
| | | Sex | 2.9667 | 1 | 0.08499 |
| | | Group | 1.6109 | 3 | 0.6569 |
| | | Day | 4.3996 | 1 | 0.03595 |
| Criterion attainment | Logistic regression | Group x Sex | -2.2746 | 3 | 0.5174 |
| | | Group | -3.2887 | 3 | 0.3492 |
| | | Sex | 4.0677 | 1 | 0.04371 |

Table S3. Statistical analysis results from experiments displayed in supplement.

| Figure | Variable | Statistical Test | Comparison | Output | *p* value |
|---|---|---|---|---|---|
| S1 | FED3, FR3 Nosepokes | rmANOVA | Group | F(3,13)=0.2317 | 0.8726 |
| | FED3, PR, breakpoint | ANOVA | Group | F(3,15)= 0.1878 | 0.9031 |
| Table S1 | Shock, flinch | ANOVA | Group | *F*(3,72)=.792 | .502 |
| | Shock, Escape | ANOVA | Group | *F*(3,72)=1.669 | .181 |
| | Shock, vocalize | ANOVA | Group | *F*(3,72)=.983 | .406 |
| Table S2 | Head sniffing time | Kruskal-Wallis | Group | *H*(3)=1.389 | .708 |
| | Head sniffing frequency | Kruskal-Wallis | Group | *H*(3)=3.960 | .266 |
| | Anogenital sniffing time | Kruskal-Wallis | Group | *H*(3)=1.386 | .709 |
| | Anogenital sniffing frequency | Kruskal-Wallis | Group | *H*(3)=2.043 | .563 |
| | Body contact time | Kruskal-Wallis | Group | *H*(3)=.810 | .847 |
| | Body contact frequency | Kruskal-Wallis | Group | *H*(3)=1.104 | .776 |
| | Total social exploration time | Kruskal-Wallis | Group | *H*(3)=1.638 | .651 |
| S2 | Tube Test, number of bouts won | ANOVA | Short, Drug | *F*(1,39)=.664 | .421 |
| | | ANOVA | Long, Drug | *F*(1,37)=.011 | .915 |
| | Social operant, FR1 mean rewards | ANOVA | Group | *F*(3,71)=.976 | .409 |
| | Day to reach learning criteria | ANOVA | Group | *F*(3,63)=2.521 | .067 |
| | Number of mice reached criteria | Fisher's Exact Test | Short, Drug | OR=.750, 95% CI[1.169,3.333] | 1.00 |
| | | | Long, Drug | OR=3.214, 95% CI[.541,19.105] | .235 |
| | FR3 rewards | rmANOVA | Group | *F*(3,55)=.425 | .736 |
| | PR3 Breakpoint | ANOVA | Group | *F*(3,75)=1.375 | .257 |
| S3 | Barnes maze, Acq distance | rmANOVA | Group | *F*(3,72)=.953 | .420 |
| | Barnes maze, Acq time | rmANOVA | Group | *F*(3,72)=.739 | .532 |
| | Barnes maze, Acq total errors | rmANOVA | Group | *F*(3,72)=3.050 | .034 |
| | Probe goal zone entries | ANOVA | Group | *F*(3,72)=.928 | .432 |
| | Probe goal zone time | ANOVA | Group | *F*(3,72)=..074 | .974 |

Table S4. Summary of current at behavior for shock sensitivity for males and females across all groups.

| Sex | Group | Flinch (mA) | Escape (mA) | Vocalization (mA) |
|---|---|---|---|---|
| Females | Short Veh | 0.17 ± 0.01 | 0.28 ± 0.03 | 0.27 ± 0.02 |
| | Short Oxy | 0.17 ± 0.01 | 0.27 ± 0.03 | 2.23 ± 1.98 |
| | Long Veh | 0.18 ± 0.01 | 0.32 ± 0.02 | 0.27 ± 0.01 |
| | Long Oxy | 0.18 ± 0.01 | 0.3 ± 0.03 | 0.26 ± 0.01 |
| Males | Short Veh | 0.18 ± 0.01 | 0.41 ± 0.03 | 0.27 ± 0.01 |
| | Short Oxy | 0.2 ± 0.01 | 0.32 ± 0.02 | 0.27 ± 0.01 |
| | Long Veh | 0.19 ± 0.01 | 0.35 ± 0.03 | 0.28 ± 0.01 |
| | Long Oxy | 0.2 ± 0.01 | 0.3 ± 0.02 | 0.25 ± 0.02 |

Results are mean ± SEM. Sample size is n=10 per sex per group.

Table S5. Summary of juvenile dyadic interaction behaviors across all groups.

| Group | Head sniffing | | Anogenital sniffing | | Body contact | | Total social exploration time (s) |
|---|---|---|---|---|---|---|---|
| | Frequency | Duration (s) | Frequency | Duration (s) | Frequency | Duration (s) | |
| Short Veh (n=20) | 66.35 ± 2.25 | 120.67 ± 6.92 | 61 ± 2.6 | 129.9 ± 8.29 | 215.15 ± 11.24 | 117.3 ± 9.96 | 254.52 ± 15.05 |
| Short Oxy (n=10) | 65.65 ± 3.94 | 117.84 ± 9.39 | 57.8 ± 4.09 | 119.51 ± 9.54 | 212.05 ± 14.88 | 112.45 ± 11.16 | 241.24 ± 18.4 |
| Long Veh (n=20) | 64.4 ± 2.65 | 115.38 ± 6.65 | 58.2 ± 3.07 | 131.56 ± 7.6 | 216.8 ± 9.9 | 119.03 ± 10.65 | 251.1 ± 13.91 |
| Long Oxy (n=19) | 71.68 ± 2.82 | 125.41 ± 6.62 | 63.11 ± 2.74 | 134.73 ± 8.06 | 223.89 ± 12.96 | 124.58 ± 10.48 | 264.03 ± 14.07 |

Results are mean ± SEM.

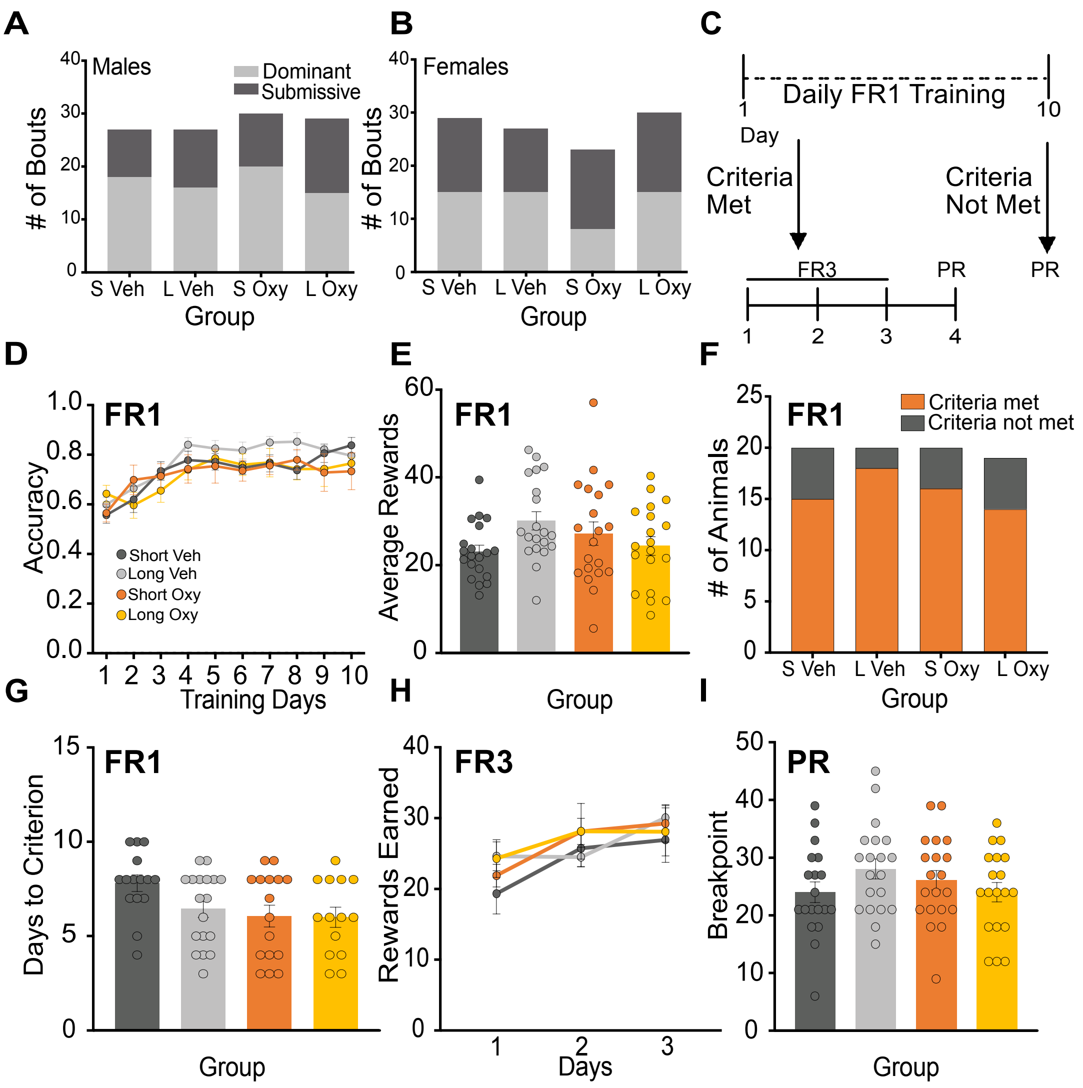


**Figure S2. Oxycodone exposure does not impact hierarchical dominance or social motivation behaviors. A-B.** Oxy-exposure did not impact number of dominant versus submissive demonstrations in the tube test in males (**A**) or females (**B**). **C.** Social operant conditioning timeline. **D.** Nosepoke accuracy as measured by correct/total nosepokes across FR1 training days were comparable across groups. **E.** Average rewards earned over the FR1 training period across treatment groups were not different. **F.** No differences were observed for number of mice to reach training criteria across groups. **G.** The average day at which criteria was met was comparable across groups. **H.** The number of rewards earned during FR3 testing was not different between groups. **I.** Progressive ratio breakpoints were comparable across all treatment groups. Grouped data are presented as means ± SEM with individual data points as filled circles.

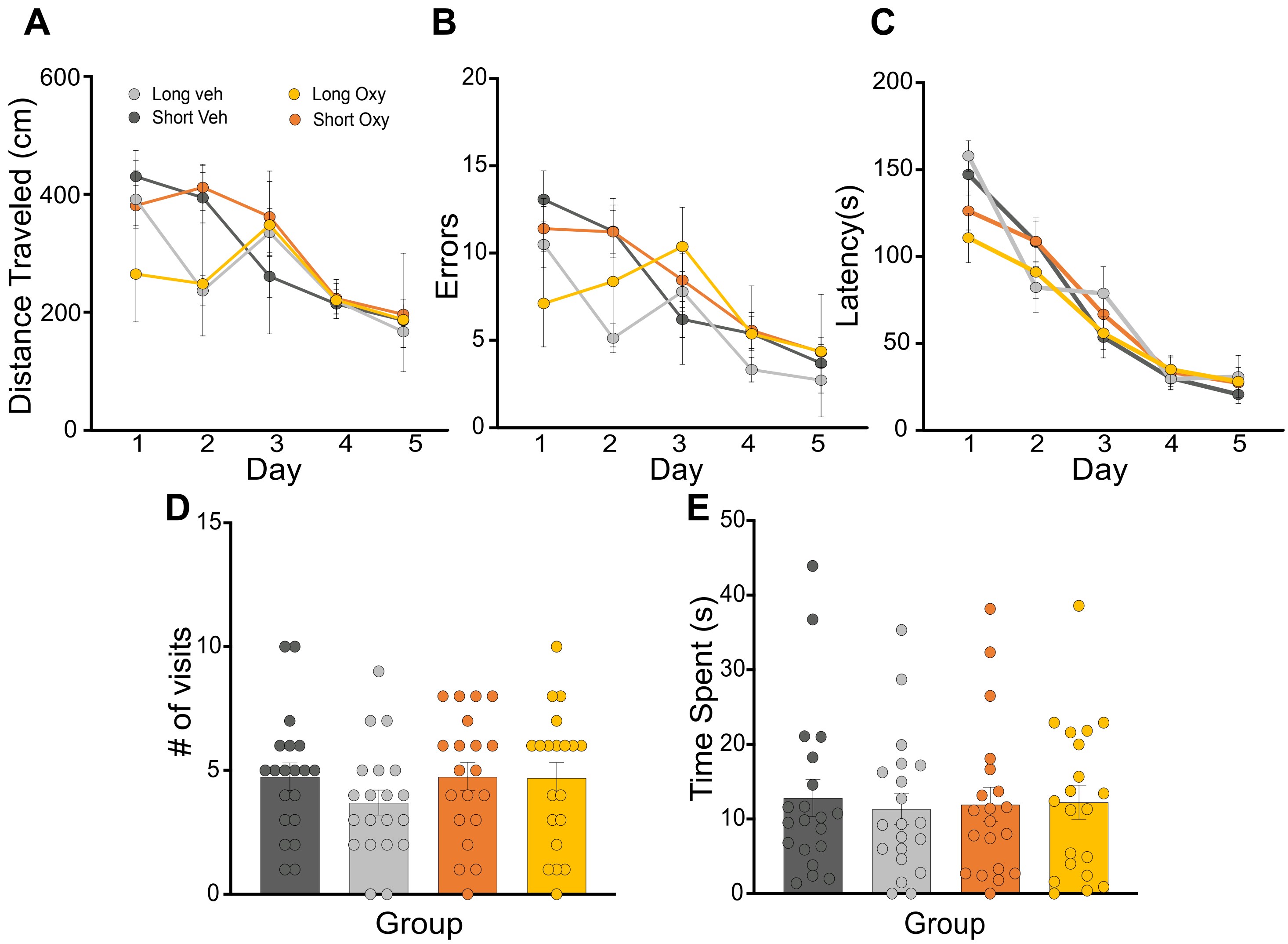


**Figure S3. Spatial memory remained intact in Oxy-exposed mice. A-C** There were no differences observed during acquisitional learning of the Barnes Maze in the distance traveled (**A**), total number of errors (**B**), and in the latency to reach the goal zone (**C**). **D-E.** There were no differences between groups in the total number of visits to the goal zone (**D**) or the time spent (**E**) in the goal zone during the probe day of the Barnes Maze.
